# The C-terminal non-catalytic domain of Mkp-1 phosphatase harbors a complex signal for rapid proteasome degradation in enterocytes

**DOI:** 10.1101/677369

**Authors:** Jin Wang, Anatoly Grishin, Patrick T. Delaplain, Christopher P. Gayer, Henri R. Ford

## Abstract

Mitogen-activated kinase phosphatase Mkp-1 is an essential negative regulator of innate immune responses. In enterocytes, Mkp-1, which is transiently expressed in response to Toll-like receptor (TLR) ligands, plays a role as an initial step in establishing tolerance to bacteria and their pro-inflammatory molecules. Mkp-1 has a very short half-life (about 30 min) in IEC-6 enterocytes. Here we examined the mechanism of Mkp-1 rapid degradation in IEC-6 cells. Immunoprecipitation and proteolysis inhibitor analysis indicated that Mkp-1 is monoubiquitinated and degraded by proteasome. Dominant negative ubiquitin increased Mkp-1 half-life, indicating involvement of ubiquitination in Mkp-1 degradation. Stability of Mkp-1 is not affected by LPS treatment, consistent with constitutive degradation. U0126, a potent and selective inhibitor of extracellular response kinase (ERK) had negligible effect on the stability of both intrinsic and ectopically expressed Mkp-1, therefore enterocytes, unlike other cell types, do not regulate Mkp-1 degradation via ERK-dependent phosphorylation. C-Truncations of the C-terminal non-catalytic domain (at amino acids 306, 331, and 357), the 306-330 deletion obliterating the potential PEST domains, as well as S296A mutation, but not S323A, S358A, or S363A mutations in potential ERK phosphorylation sites dramatically stabilized Mkp-1. C-terminal fusion of the non-catalytic C-terminal domain of Mkp-1 conferred accelerated degradation on the intrinsically stable green fluorescent protein. According to these results, rapid degradation of Mkp-1 in IEC-6 enterocytes is ubiquitin-proteasome-dependent and ERK-independent. Multiple elements of the C-terminal non-catalytic domain play essential roles in the formation of the complex rapid degradation signal.

## Introduction

Mitogen-activated protein kinase (MAPK) phosphatase-1 (Mkp-1), also known as dual specificity phosphatase 1 (Dusp1) is a member of the family of MAPK phosphatases, the enzymes that deactivate MAPKs of the extracellular response kinase (ERK), c-Jun N-terminal kinase (JNK), and p38 families, by reversing their activating threonine-tyrosine phosphorylation. MAPKs and their phosphatases regulate diverse processes including, but not limited to proliferation, survival, apoptosis, development, stress response, and inflammation. Mkp-1 has a particularly prominent role as a negative regulator of inflammation. Mice deficient in this phosphatase are healthy under normal conditions, but show exaggerated inflammatory response to various pathogens^1–7^. Consistent with its role as dynamic inhibitor of inflammation, Mkp-1 has low basal levels, but is induced by inflammatory stimuli^8–11^ and then rapidly degraded via ubiquitin-proteasome, with the half-life of 1-2 h depending on the cell type^12–14^.

Mkp-1 is involved in the establishment of tolerance to bacteria in the intestinal epithelium. Naïve enterocytes are highly sensitive to Toll-like receptor (TLR) ligands, bacterial components including cell wall macromolecules, flagellin, dsRNA, and unmethylated CpG DNA, which trigger innate immune responses mediated by MAPKs and the pro-inflammatory transcription factor NF-κB^15, 16^. The latter mediates rapid induction of Mkp-1, which transiently blunts MAPK-dependent inflammatory responses^16^ before profound and long-lasting shutdown of TLR signaling and establishing tolerance to bacteria via induction of the highly stable ubiquitin-editing enzyme A20^17^.

Proteasome-dependent degradation of Mkp-1 depends on its C-terminal non-catalytic domain, a hydrophilic sequence rich in proline, glutamic acid, serine, and threonine (PEST) residues. Within this domain, four ERK phosphorylation sites have been identified as signals affecting the rate of proteasome-dependent proteolysis^12–14, 18^. Otherwise, structures within the non-catalytic domain that serve as degradation signal(s) remained unknown.

Here we report very short (30 min) half-life of Mkp-1 in enterocyte cell lines due to ubiquitin-proteasome degradation. We demonstrate that rapid degradation of Mkp-1 in IEC-6 enterocytes is constitutive and ERK-independent, and that multiple elements of the C-terminal non-catalytic domain play essential roles in the formation of the rapid degradation signal.

## Materials and Methods

### Reagents

FLAG M2 and β-actin antibodies, proteasome and protease inhibitors, U0126, LPS from *E. coli* 0127:B8 were purchased from Sigma-Aldrich, St. Louis, MO. Poly (IC) was from Pharmacia, Uppsala, Sweden. GFP B2, Mkp-1 M18, ubiquitin P4D1 were from Santa Cruz Biotechnology, Santa Cruz, CA. Phospho-ERK antibody was from Cell Signaling Technology, Danvers, MA.

### Cell culture

IEC-6, IEC-18 (ATCC, Manassas, VA) and RIE (a gift of Dr. Pawel Kiela, University of Arizona, Tucson, AZ) cells at passages 15-30 were grown in Dulbecco-modified Eagle medium (DMEM) supplemented with 5% fetal bovine serum (FBS) at 37°C and 10% CO_2_. SW480 cells (ATCC) were grown in Leibowitz’s L15 medium with 10% FBS at 37°C in air, without CO_2_ Cultures at 70-90% confluence were used in experiments.

### Plasmids

pFLAG-MKP1 has been described previously^16^. pFLAG-GFP was constructed by PCR-amplifying GFP coding sequence from pMAX-GFP (Lonza, Alpharetta, GA) with GGGAATTCAAGTAAAGGAGAAGAACTTTTC and GGGATCCCTATTTGTATAGTTCATCCATGCC primers, cutting the resulting PCR product with *Eco*RI and *Bam*HI, and inserting it into *Eco*RI- and *Bam*HI-cut p3xFLAG-CMV10 (Sigma-Aldrich). The GFP-Ub K0.G76V zero lysine GFP-ubiquitin fusion plasmid^19^ was obtained from Addgene, Watertown, MA.

### In vitro mutagenesis

Mutations were introduced into plasmid constructs using the QuickChange Site-Directed Mutagenesis Kit (Agilent Technologies, San Jose, CA) as recommended by the manufacturer. Mutation sites and adjacent regions were sequenced to verify absence of errors.

### Real time RT-PCR

Total cellular RNA was purified using Trizol (Thermo Fisher Scientific, Canoga Park, CA) as recommended by the manufacturer. RNA concentrations were measured on NanoDrop (Agilent Technologies). First strand synthesis and real time PCR were performed using iScript kit and iTaq Universal SYBR Green Supermix (Bio-Rad, Hercules, CA) on Light Cycler 480 (Roche, Indianapolis, IN). The following primers were used: CTGAGTACTAGTGTGCCTGACAGT and TTTCCGGGAAGCATGGTAAGCACT (MKP-1); GTTCGCACCAAGACTGTGAAGAAG and ATCCGCTTCATCAGATGCGTGACA (RPS17). Levels of MKP-1 transcript were normalized to those of RPS17 mRNA in the same sample. Melting curve analysis of PCR products was performed to ascertain the absence of genomic sequence amplification.

### Immunoprecipitation

IEC-6 cells were treated with 0.1 μg/ml LPS + 3 μM MG-132 for 30 min. Cells were lysed on ice with RIPA buffer (50 mM Tris-HCl pH 7.5, 100 mM NaCl, 1% NP40, 0.5% sodium deoxycholate, 0.1% SDS, 1mM PMSF) for 10 min. After clearing by centrifugation at 10,000xg for 10 min, lysates were incubated with 1 μg/ml Mkp-1 antibody for 1 h at 4°C. Antibody-antigen complexes were collected on protein A sepharose, washed 3 times with RIPA buffer, and analyzed by Western blotting.

### Amino acid sequence analysis

Potential PEST domains were identified and scored using the algorithm epestfind at emboss.bioinformatics.nl

### Western blots, immunofluorescence microscopy

were performed as previously described^16^.

### Statistical analysis

Standard deviations were determined using GraphPad (www.graphpad.com).

## Results and Discussion

### Mkp-1 is rapidly degraded in enterocytes via proteasome-dependent pathway

Mkp-1 induction is an early step in desensitization of intestinal epithelial cells to pro-inflammatory TLR ligands^16^. TLR ligand-induced expression of Mkp-1 in IEC-6 enterocytes of rat origin is transient: Mkp-1 protein levels peak at 30-60 min and return to baseline in 120 min following treatment with dsRNA and flagellin, the ligands of TLR3 and TLR5, respectively (Fig. 1A), as well as LPS and unmethylated CpG DNA, the ligands of TLR4 and TLR9^16^. Similar results were obtained using two other primary rat enterocyte cell lines, IEC-18 and RIE, as well as human colonic cell line SW480 (data nor shown). Since transient expression of Mkp-1 may be important for dynamic regulation of innate immune response in the intestinal epithelium, we sought to gain insight into the underlying mechanisms. First, we asked whether the transient expression is a property of TLR-mediated induction, or a general characteristic of Mkp-1. In addition to TLR ligands, Mkp-1 is induced by a variety of stresses in IEC-6 cells, but most stresses cause relatively weak induction (data not shown), which complicates degradation measurements. We have found that hypoxic stress (5% O_2_) causes relatively strong upregulation of Mkp-1, which steadily increases within 120 min of treatment (Fig. 1A). Differences in the time course of Mkp-1 expression could be due to different mRNA levels and/or speed of protein degradation. To test the first possibility, we measured MKP-1 transcript levels at several time points of treatment with dsRNA, flagellin, and hypoxia using real time RT-PCR. Whereas TLR ligands caused transient induction of MKP-1 mRNA peaking at 30-60 min, hypoxia caused steady accumulation of this mRNA (Fig. 1B). In these experiments, MKP-1 mRNA levels closely followed Mkp-1 protein expression, which is consistent with rapid degradation of the Mkp-1 protein under both treatment scenarios.

**Fig. 1.**
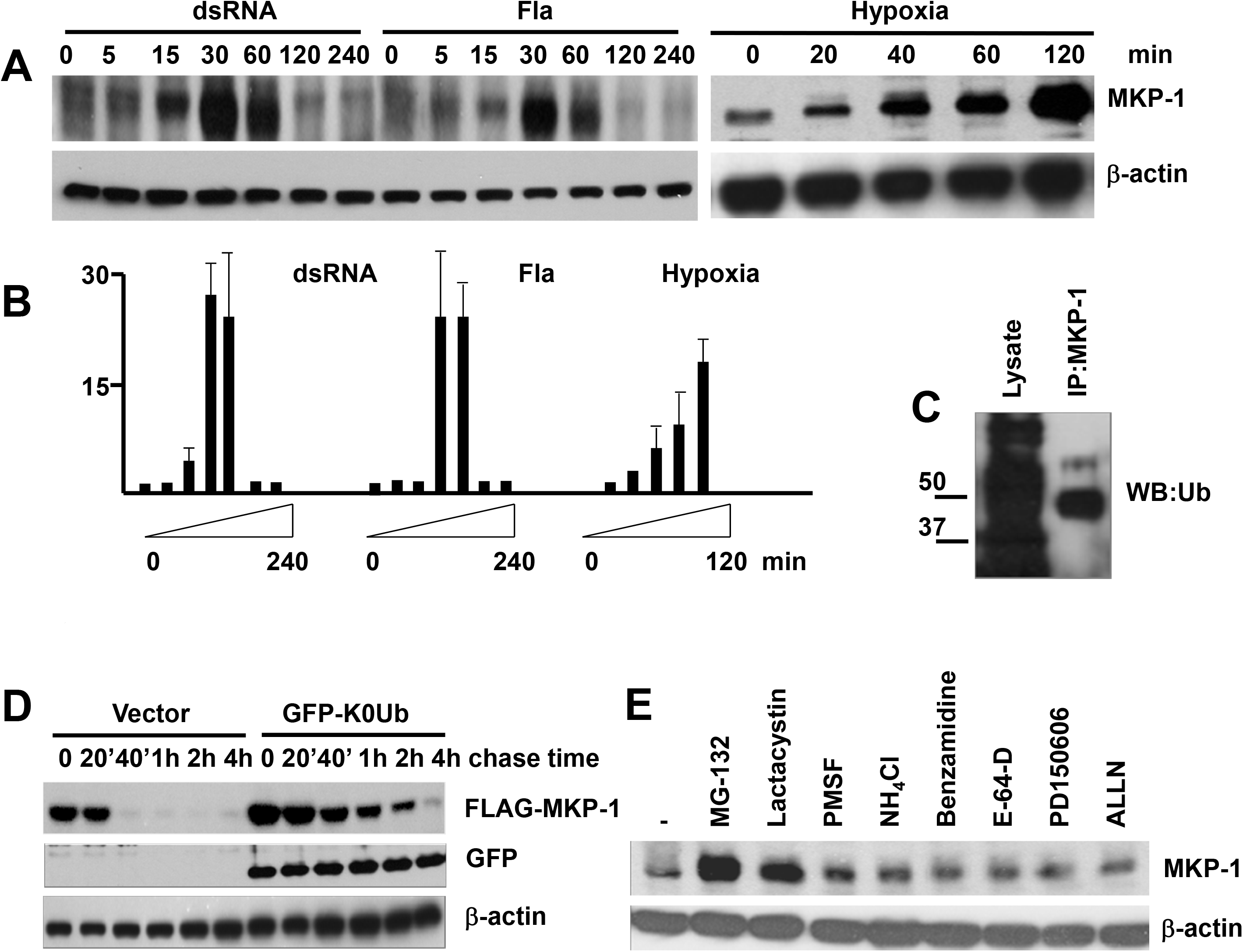
Time course and proteasome dependency of Mkp-1 degradation. A, Mkp-1 protein levels in IEC-6 cells treated with 50 μg/ml dsRNA, 0.1 μg/ml flagellin, or hypoxia (5% 0_2_) for indicated time. B, Mkp-1 transcript levels in IEC-6 cells treated as in A, average of 4 independent experiments. C, Mkp-1 ubiquitination. IEC-6 cells were treated with 0.1 μg/ml LPS + 3 μM MG-132 for 60 min. Cell lysate was immunoprecipitated with Mkp-1 antibody and immunoprecipitate analyzed by Western blotting for ubiquitin. Positions of molecular size markers (kD) are indicated on the left. D, cycloheximide chase assay for FLAG-MKP-1 stability in cells co-transfected with pFLAG-MKP1 and either empty vector or GFP-Ub K0.G76V. E, effects of protease inhibitors on MKP-1 levels in IEC-6 cells. Cells were treated with LPS + 3 μM MG-132, 10 μM lactacystin, 1 mM PMSF, 10 mM NH_4_Cl, 1 mM benzamidine, 10 μM E-64-D, 10 μM PD150606, or 10 μM ALLN as indicated, for 2 h. Western blot images are representative of at least 3 independent experiments.

Previous studies implicated ubiquitin-proteasome in rapid degradation of Mkp-1^12–14, 20–25^. To test whether ubiquitin is involved in Mkp-1 degradation in enterocytes, we immunoprecipitated Mkp-1 from IEC-6 cells treated with LPS and probed the resulting immunoprecipitate for conjugated ubiquitin by Western blotting. There was a prominent band whose electromobility was consistent with monoubiquitinated Mkp-1 species (Fig. 1B). Monoubiquitination of Mkp-1 was also reported by others^13^. To examine functional role of ubiquitin in Mkp-1 degradation, we co-transfected IEC-6 cells with FLAG epitope-tagged MKP1 and dominant negative (zero lysine) ubiquitin constructs, and examined stability of Mkp-1 using the cycloheximide chase assay. Dominant negative ubiquitin increased the half-life of FLAG-Mkp-1 from 30 min to 1 h (Figure 1D), indicating participation of ubiquitin in Mkp-1 degradation. To elucidate the role of proteasome, we determined the levels of Mkp-1 in IEC-6 cells treated with LPS in the presence of various proteolytic inhibitors. Specific inhibitors of proteasome, MG-132 and lactacystin, but not inhibitors of serine proteases (PMSF, benzamidine), lysosomal proteases (NH_4_Cl, E-64-D), calpain (PD150606) or cathepsin (ALLN), resulted in accumulation of Mkp-1 (Fig. 1E), pointing to the specific role of proteasome. Thus, enterocytes, like other cell types, utilize the ubiquitin-proteasome system to rapidly degrade Mkp-1.

### Degradation of Mkp-1 in enterocytes is not affected by LPS

TLR ligand-induced proteolysis might explain transient expression of Mkp-1 in TLR ligand-treated, as opposed to sustained expression in hypoxia-treated enterocytes. To test the possibility of TLR-mediated degradation, we transfected IEC-6 cells with FLAG-MKP-1, treated the transfectants with LPS, and examined Mkp-1 levels at various times of LPS treatment. Levels of Mkp-1 were not affected by LPS (Fig. 2A). Moreover, the half-life of FLAG-Mkp-1 (about 30 min as measured by the cycloheximide chase assay) was not affected by LPS (Fig. 2B). Therefore, rapid degradation of Mkp-1 in enterocytes does not depend on an LPS-regulated proteolytic system.

**Fig. 2.**
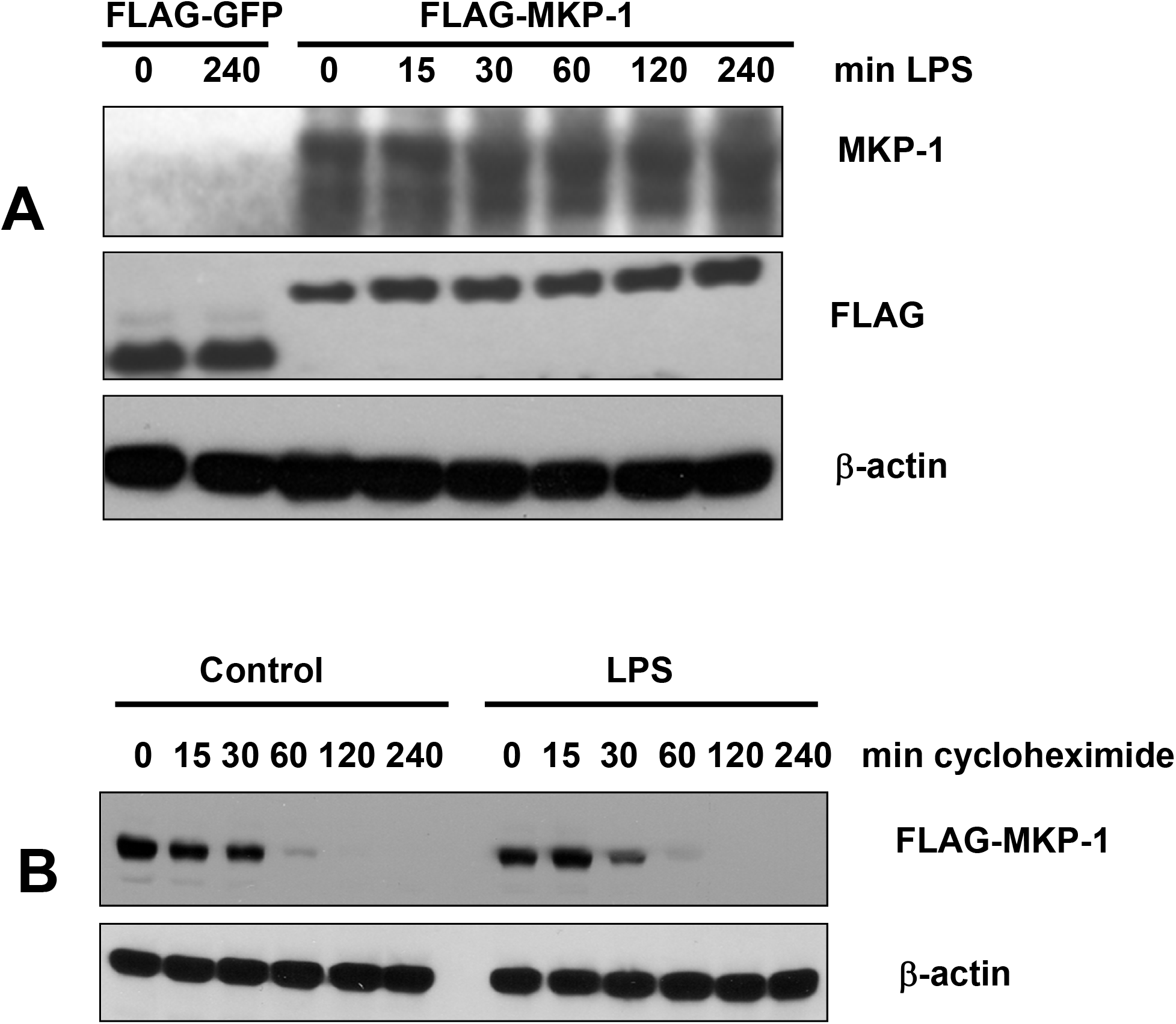
LPS does not affect levels of ectopically expressed Mkp-1. A, IEC-6 cells transiently transfected with pFLAG-GFP or pFLAG-MKP-1 were treated with 0.1 μg/ml LPS for indicated times. Levels of Mkp-1, FLAG, and β-actin were examined by Western blotting. B, IEC-6 cells transiently transfected with pFLAG-MKP1 were treated with 10 μM cycloheximide for indicated times in the absence (control) or presence of LPS, and levels of FLAG-tagged Mkp-1 and β-actin were examined. Images are representative of at least 3 independent experiments.

### Degradation of Mkp-1 in enterocyte is not regulated by ERK

Previous studies in non-enterocyte cell lines implicated ERK-dependent phosphorylation at the non-catalytic C-terminal domain of Mkp-1 as a modulator of its degradation. As some studies found ERK phosphorylation accelerating^14, 18, 20, 21^, and others delaying Mkp-1 degradation^12, 13, 23, 24, 26, 27^, the role of phosphorylation remains controversial. In the epithelium of rat small intestine, both Mkp-1 and activated ERK localize to crypts (Fig. 3A), raising the possibility of functional interaction between these two proteins. However, ERK blockade with U0126, a potent and highly selective inhibitor of ERK phosphorylation, did not appreciably alter the time course of LPS-induced Mkp-1 expression (Fig. 3B). In addition, U0126 had negligible effect on the half-life of Mkp-1 (Fig. 3C). Accordingly, ERK plays minimal, if any, role in Mkp-1 degradation in enterocytes. Because Mkp-1 half-life (about 30 min) remained constant under different scenarios, the rapid proteasome-dependent degradation of this MAPK phosphatase in enterocytes is likely constitutive.

**Fig. 3.**
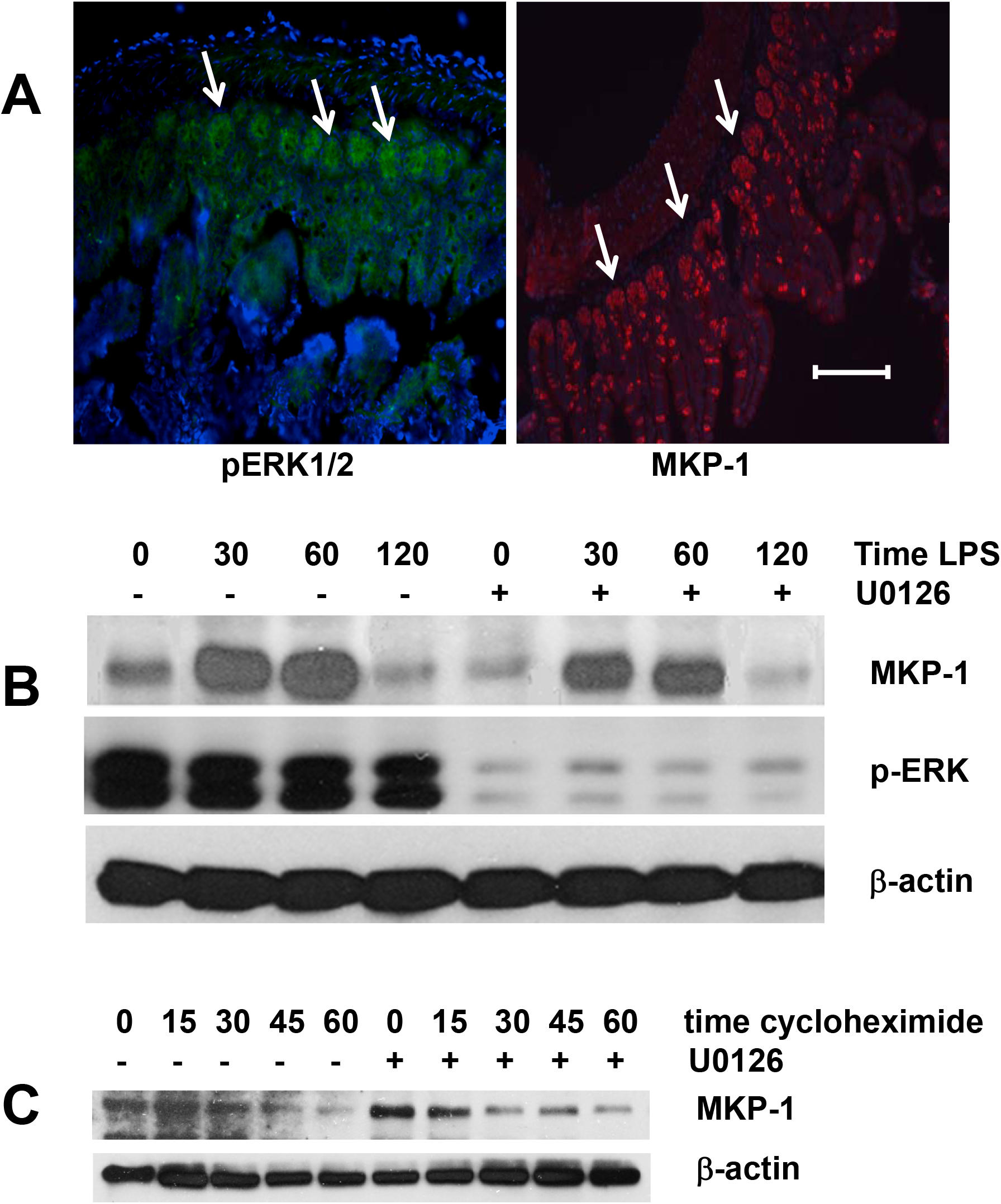
Effect of ERK on Mkp-1 expression. A, phospho-ERK (green) and Mkp-1 (red) immunofluorescence in rat ileum. Arrows point to intestinal crypts. Blue, DAPI-stained nuclei. Bar=100 μm. Images are representative of samples from 4 animals. B, time course of LPS-induced Mkp-1 expression in IEC-6 cells in absence or presence of 2 μM ERK phosphorylation inhibitor U0126. C, time-dependent degradation of Mkp-1. IEC-6 cells were treated with LPS for 30 min, without or with U0126, and chase was started by adding cycloheximide to 10 μM. Western blots are representative of at least 3 independent experiments.

### The C-terminal non-catalytic domain of Mkp-1 harbors a complex signal for rapid degradation

Although the C-terminal non-catalytic domain of Mkp-1 is rich in PEST amino acids (Fig. 4A), the potential PEST sequences within have negative PEST scores (Fig. 4A), disqualifying them as classical PEST domains^28^. In order to identify structures responsible for the rapid degradation of Mkp-1, we performed in vitro mutagenesis of its C-terminus and determined half-lives of the resulting mutants using the cycloheximide chase assay. Truncations at residues 307, 331, and 357 all dramatically stabilized Mkp-1, increasing its half-life from 30 min to over 240 min (Fig. 4B, C). The 307-330 deletion obliterating the two potential PEST sequences had the same effect (Fig. 4C). Point mutations substituting serine for alanine in potential ERK phosphorylation sites (SP) at positions 323, 358, and 363 had no effect on Mkp-1 stability, whereas alanine at 296 caused dramatic stabilization. Results of mutagenesis experiments identify S296, potential PEST sequences, and the C-terminal sequence SPITTSPSC as essential parts of a complex rapid degradation signal. To test whether the C-terminal non-catalytic domain of MKP-1 is sufficient to direct protein degradation, we fused the last 78 amino acids of Mkp-1 to the C-terminus of an intrinsically stable green fluorescent protein (GFP) and examined the effect of the Mkp-1 adduct on GFP stability. The C-terminus of Mkp-1 decreased the half-life of GFP from over 12 h to 4 h, indicating that it was capable of conferring increased rate of degradation on an intrinsically stable protein. The fact that GFP-Mkp-1_C_ fusion degraded at slower rate compared to Mkp-1 may indicate that sequences outside of the non-catalytic domain of the latter enhance the rapid degradation signal, or that GFP moiety exerts an inhibitory effect on degradation.

**Fig. 4.**
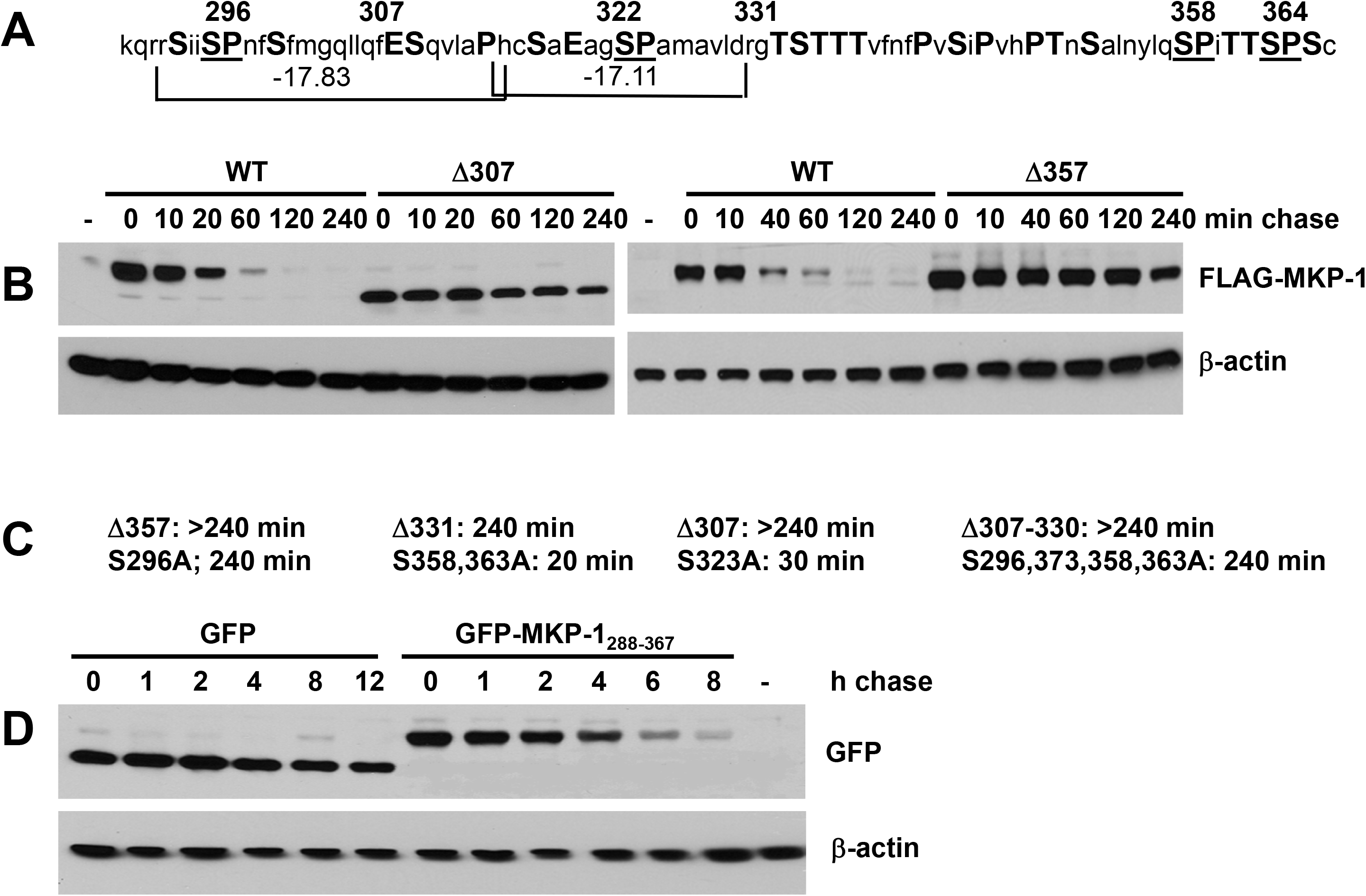
The C-terminal non-catalytic domain of Mkp-1 harbors a complex degradation signal. A, amino acid sequence of the C-terminal non-catalytic domain in single letter code. PEST residues are capitalized. Potential ERK phosphorylation sites (SP) are underscored. Numbers on top indicate amino acid positions beginning from the N-terminus. Brackets at the bottom delineate potential PEST domains, with their PEST scores shown. B, cycloheximide chase assay for stability of WT Mkp-1 and C-terminal truncations at 307 and 357 expressed in IEC-6 cells as FLAG-tagged proteins. C, half-lives of the indicated MKP-1 mutants. D, cycloheximide chase assay for stability of WT Green Fluorescent Protein (GFP) and GFP fused at the C-terminus to the non-catalytic C-terminal domain of Mkp-1. -, cells transfected with empty vector. Western blot images are representative of at least 3 independent experiments.

Taken together, our results indicate that rapid ubiquitin-proteasome-dependent degradation of MKP-1 in enterocytes is largely similar to that in other cell types, with exceptions for speed and the role of ERK phosphorylation. Site-directed mutagenesis of ERK phosphorylation sites has been previously used as a tool to probe the role of ERK in Mkp-1 degradation^12–14^. Our results demonstrate that such mutations, in addition to their intended effect, may affect degradation by perturbing the basic constitutive C-terminal degradation signal, regardless of ERK phosphorylation. It is also plausible that enterocytes are lacking the auxiliary proteasomal degradation factors that recognize phosphorylated serine residues within the C-terminal non-catalytic domain of Mkp-1. This study reveals the complex character of the Mkp-1 degradation signal and shows that strong protein degradation domains may not necessarily conform to the PEST hypothesis.

## Acknowledgments

We would like to thank Nico Dantuma for the opportunity to use the GFP-Ub K0.G76V plasmid, and Paul Kiela for the gift of RIE-1 cells. This study was supported by NIH Grant AI 014032 to HRF.

